# Sigmoni: classification of nanopore signal with a compressed pangenome index

**DOI:** 10.1101/2023.08.15.553308

**Authors:** Vikram S. Shivakumar, Omar Y. Ahmed, Sam Kovaka, Mohsen Zakeri, Ben Langmead

**Affiliations:** Department of Computer Science, Johns Hopkins University

## Abstract

Improvements in nanopore sequencing necessitate efficient classification methods, including pre-filtering and adaptive sampling algorithms that enrich for reads of interest. Signal-based approaches circumvent the computational bottleneck of basecalling. But past methods for signal-based classification do not scale efficiently to large, repetitive references like pangenomes, limiting their utility to partial references or individual genomes. We introduce Sigmoni: a rapid, multiclass classification method based on the *r*-index that scales to references of hundreds of Gbps. Sigmoni quantizes nanopore signal into a discrete alphabet of picoamp ranges. It performs rapid, approximate matching using matching statistics, classifying reads based on distributions of picoamp matching statistics and co-linearity statistics. Sigmoni is 10-100*×* faster than previous methods for adaptive sampling in host depletion experiments with improved accuracy, and can query reads against large microbial or human pangenomes.

## 1 Introduction

Read classification is the core of many sequencing data analyses, including taxonomic classification [1]–[3], host depletion [4], and adaptive sampling [5]–[7]. In all these applications, the completeness of the reference sequences used can impact classification accuracy. Incomplete references can cause misclassification of reads with novel variation [4]. Compressed full-text indexes like *r*-index can index larger collections containing more variation, effectively avoiding reference bias while also coping with the increasing size of reference-genome databases [8], [9].

The computational bottleneck for nanopore read classification is basecalling. Early nanopore basecallers were based on Hidden Markov Models (HMMs), which were efficient but yielded base-level accuracy as low as 70% [10]. Recent basecallers like Guppy [11] use recurrent neural networks (RNNs), achieving more than 90% base-level accuracy. However, these networks comprise millions of parameters and are computationally infeasible to run without hardware acceleration [12]. Even using acceleration, basecalling is the rate-limiting step for read classification algorithms [13]. One study found that over 95% of the compute time in a variant calling pipeline for SARS-CoV2 was spent in basecalling [14].

Still, many base-level methods for adaptive sampling have been proposed; Readfish [6] uses Minimap2 [15] to align basecalled reads and perform adaptive sampling. SPUMONI [4] uses the *r*-index to query basecalled reads against large collections of references efficiently. SPUMONI 2 [16] decreases index size and improves speed using minimizer digestion. SPUMONI 2 also introduces a document array data structure for multi-class classification. TargetCall [13] introduced the idea of pre-basecalling filtering, using simpler neural-network basecallers to identify reads of interest, which are then re-basecalled with larger, slower models. All of these methods rely on basecalling, and therefore do not address the main bottleneck.

Signal-based nanopore read classification methods have also been proposed. UNCALLED [5] probabilistically identifies *k*-mer sequences from the nanopore signal, while aligning the *k*-mers to an FM-index and clustering co-linear matches to determine read origin. However, UNCALLED becomes infeasible for large references (*>* 1 Gbp) because the FM-index scales linearly with reference length. Also, the seed clustering process requires superlinear time when matching to repetitive references. Another method, Sigmap, [7] queries the read signal against a *k-d* tree built over the expected signal from the reference. Sigmap chains candidate seeds using dynamic programming. However, the *k-d* tree suffers from poor scaling for large references and the seed-chaining method becomes inefficient for repetitive reads with many candidate hits. Lastly, RawHash [17] performs a similar seed-chain-extend method using a hash table over the reference sequence to identify seed matches, but has similar drawbacks as Sigmap and UNCALLED.

Other signal-based methods are application-specific. RawMap [18] uses support vector machines to identify signal patterns corresponding to microbial species in order to filter non-human reads. SquiggleNet [19] uses a convolutional neural network to distinguish human from bacterial reads. However, these methods do not generalize to different classification tasks.

Here, we introduce Sigmoni, which extends the *r*-index framework for read classification – first used in SPUMONI – to the problem of classifying raw nanopore electrical signal. Sigmoni uses an ultra-fast signal discretization method to project the current signal into a small alphabet for exact match querying with the *r*-index. With a rapid pre-processing and querying step, along with efficient reference indexing, Sigmoni can classify reads faster than previous signal-based classification methods against complete reference sequences for accurate and rapid filtering and adaptive sampling.

## 2 Results

### 2.1 Method overview

Sigmoni classifies reads by identifying stretches of signal that approximately match a reference sequence. The sequencer emits regular picoamp measurements, which Sigmoni segments and quantizes into a sequence of symbols from a discrete alphabet of picoamp ranges. By default, Sigmoni uses an alphabet of size 6, which we found balanced robustness to noise with specifity in exact matching. Prior to indexing, reference sequences are also converted to the picoamp alphabet using a “pore model” that maps *k*-mers to their expected picoamp level. The reference index will include sequences from two or more classes. These might represent a positive (on-target) and null (off-target) class, or they might represent multiple classes, e.g. for distinguishing species in a metagenomics sample.

Sigmoni, like SPUMONI, uses matching statistics and the related idea of pseudo-matching lengths (PMLs) to make classification decisions. When querying with a length-*n* read, the length-*n* vector of matching statistics encode the length of the longest half-maximal-exact-match (half-MEM) extending from each query position. The statistics are large in portions of the query that match the reference well. In areas that do not match well, they will be short, representing “random” coincidental matches. Classification decisions can be made based on the PML distribution with respect to sequences in the index. E.g. SPUMONI used a KS-test to ask if matching statistics were longer with respect to the positive class compared to the null class.

PMLs are similar to matching statistics, with both being effective for classification problems [4]. Sigmoni uses PMLs since they can be computed using a single fast loop over the input, whereas matching statistics require either multiple loops (as in MONI [20]) or more complex loops (as in PHONI [21]).

To co-localize matches to the reference, Sigmoni partitions the reference sequences into equal-length sections called “shreds.” Using the sampled document array proposed in SPUMONI 2 [16], a given match (e.g. a peak in the PML vector) can be associated to a shred. PML-shred associations are aggregated to identify whether matches tend to co-localize in a particular shred, analogous to the seed clustering or chaining step in a read aligner. Additionally, these matches are weighted by their sequence complexity to down-weight matches to repetitive regions. A final classification decision is made by assigning the read to the reference sequence containing the shred that received the most (weighed) matches.

### 2.2 Binary classification

#### Mock community

To evaluate Sigmoni’s binary classifications, we used a dataset of reads previously sequenced [5] from the ZymoBIOMICS High Molecular Weight DNA Mock Microbial community (Zymo). This consists of one yeast species (*Saccharomyces cerevisiae*) and seven bacterial species (*Staphylococcus aureus, Salmonella enterica, Escherichia coli, Pseudomonas aeruginosa, Listeria monocytogenes, Enterococcus faecalis*, and *Bacillus subtilis*). The task was to distinguish yeast-origin reads (comprising ∼1% of the dataset) from bacterial reads. We used the F1 metric, the harmonic mean of precision and recall, with read-length sample weighting (prioritizing correctly classifying longer reads). True read-origin labels were determined by using Minimap2 [15] to align basecalled versions of the reads to the mock community reference genomes, giving reads that uniquely mapped to a single reference (mapping quality ≥ 30) the corresponding label. Reads for which basecalled versions did not map were excluded from evaluation. Note that the basecalled reads are used only to obtain the truth labels; the signal-based methods are compared using the signal data (not basecalled) for the same reads. Similar to previous signal-based studies, we omitted basecalled classification methods (e.g. Readfish) from comparison as the basecalling step makes them notably inefficient [14] and thus incomparable to signal-based methods.

Overall, Sigmoni yielded higher precision and lower recall than other methods (Table 1, left). Sigmoni’s index of mock community genomes – consisting of both the *r*-index and document array built over a total of 41.2 Mbp of reference sequence – fit in 178 MB. UNCALLED, which uses an FM-index, had the smallest index size (72 MB), though its F1 was low due to low precision. RawHash, which was run with the recommended -x sensitive parameter, was unable to classify over 10% of reads, while Sigmoni classified all reads in the dataset. RawHash and Sigmap also had much larger index sizes: 567 and 2,591 MB, or 3.2-fold and 14.6-fold larger (respectively) than Sigmoni’s index.

**Table 1:** Comparison of binary classification between signal-based methods in mock community (distinguishing yeast from bacterial-origin reads, left), and host depletion (distinguishing human from mock-community reads, right) experiments.

|  | Zymo |  |  |  |  | Human (host depletion) |  |  |  |  |
| --- | --- | --- | --- | --- | --- | --- | --- | --- | --- | --- |
|  | Precision | Recall | F1 weighted | Unclassified rate | Index size (MB) | Precision | Recall | F1 weighted | Unclassified rate | Index size (GB) |
| **Sigmoni** | 1.0 | 0.928 | 0.963 | 0.0 | 177.6 | 0.882 | 0.979 | 0.929 | 0.0 | 14.7 |
| Sigmap | 0.956 | 0.949 | 0.952 | 0.046 | 2591.3 | 0.831 | 0.954 | 0.888 | 0.382 | 175.9 |
| Uncalled | 0.460 | 0.986 | 0.627 | 0.044 | 72.1 | * | * | * | * | * |
| RawHash | 0.967 | 0.941 | 0.954 | 0.1108 | 567.4 | 0.591 | 0.998 | 0.742 | 0.5126 | 13.7 |

#### Host depletion

Host depletion is a common task in metagenomics experiments, e.g. in microbiome studies where reads from the human host are less scientifically relevant and create the risk of identifying the host individual. To emulate a host depletion scenario, we created a hybrid dataset of real nanopore reads from the NA12878 WGS consortium [22] and Zymo mock community reads at equal proportions. As in the previous section, we evaluated each method’s ability to discriminate human from Zymo reads using the read length-weighted F1 metric.

Sigmoni achieved the greatest length-weighted F1, while also classifying the full read set (Table 1, right). RawHash did not classify over 50% of reads, and Sigmap did not classify 38%. Since unclassified reads were predominantly human-origin, we tallied these in the same category as reads explicitly classified as human, allowing Sigmap and RawHash to perform competitively on F1 score. UNCALLED was unable to index the human genome and mock community genomes, and was omitted from this comparison.

Sigmap’s index was 176 GB, too large to fit in physical RAM on a typical, non-server computer. RawHash and Sigmoni created indexes of similar size (14 and 15 GB respectively). In RawHash’s case, this is achieved with a minimizer digestion scheme that likely contributed to its lower classification rate. Sigmoni was able to classify the full dataset with the best length-weighted F1 (0.929), while using a small index: about the same size as RawHash’s and 12-fold smaller than Sigmap’s.

### 2.3 Multi-class classification

#### Mock community

Sigmoni can perform multi-class classification using a “sampled document array” structure to identify a reference region (“shred”) of origin for each exact match. The sampled document array was first introduced in the SPUMONI 2 study [16] and is detailed in Methods 4.5. We used the the same Zymo mock community dataset (section 2.2) to classify species of origin (one of *S. aureus, S. enterica, E. coli, P. aeruginosa, L. monocytogenes, E. faecalis, B. subtilis* or *S. cerevisiae*). We assessed multi-class classification performance using read-length weighted accuracy: the proportion of reads that received a correct label assignment, weighted by read length. For UNCALLED, Sigmap, and RawHash, we used the same output alignments as in the binary task, however, we now considered all eight species as separate classes, rather than just bacterial and yeast classes.

Figure 2**A** shows confusion matrices for each method. Diagonal entries (i.e. true positives) are omitted, and the color scale adjusted to highlight misclassified reads, represented in off-diagonal elements. Consistent with the binary classification results (Table 1, left), Sigmoni classified substantially more reads than the other tools. Sigmoni had comparable accuracy to Sigmap, with Sigmoni achieving 0.964 length-weighted accuracy and Sigmap achieving 0.969. UNCALLED misclassified many reads as yeast, likely due to low complexity regions in the eukaryotic yeast genome. This yielded a low binary classification F1, however due to the low proportion of yeast reads in the dataset, UNCALLED still performed comparable in multiclass accuracy. RawHash’s accuracy was the lowest, due to the proportion of unclassified reads.

**Figure 1:**
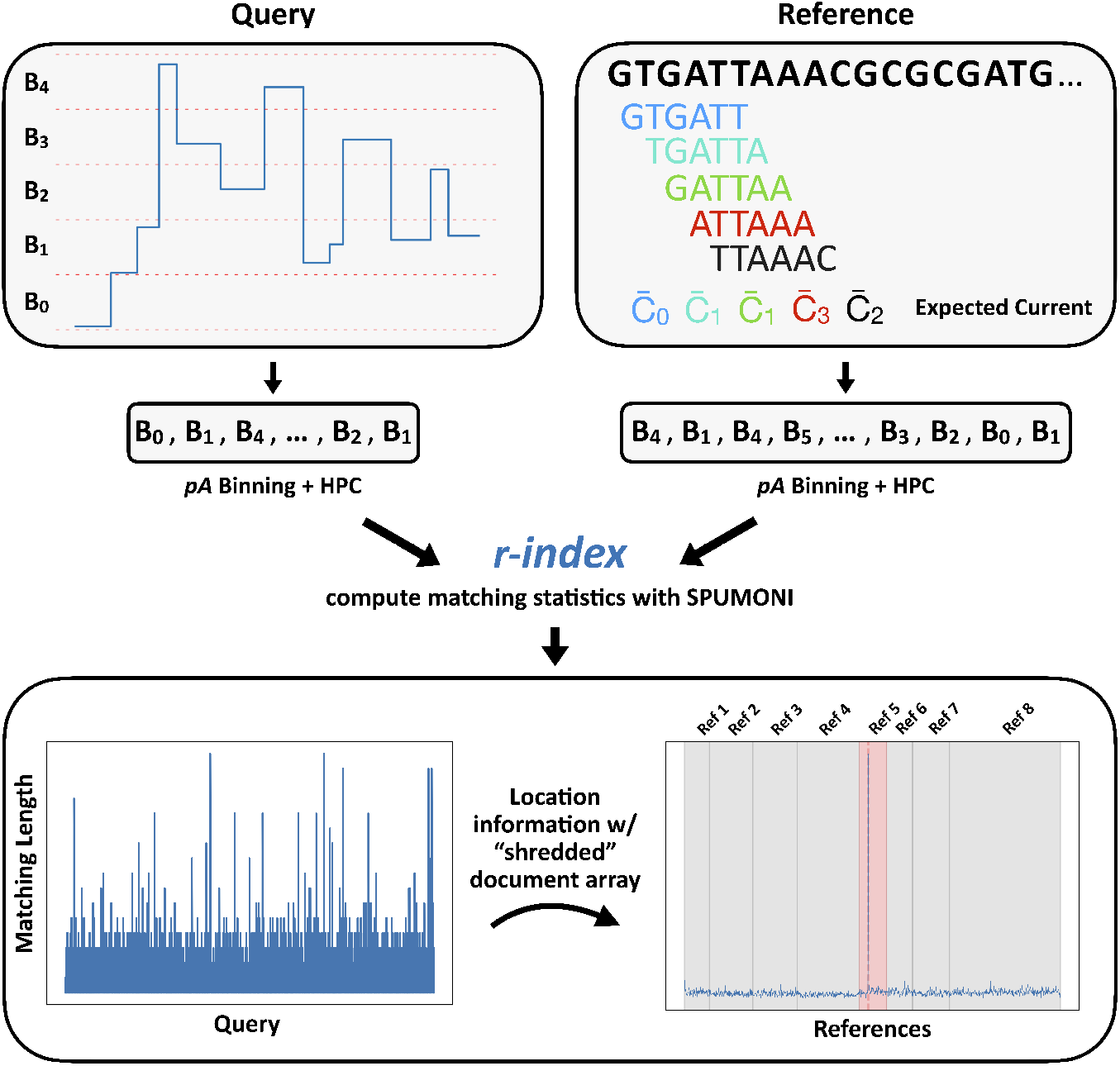
Overview of Sigmoni mapping procedure: (top) Query is discretized into “bins”, which are further converted into arbitrary characters from a small alphabet for exact matching. The reference is digested into *k*-mers and converted to the same alphabet based on the expected current level. (bottom) The matching length profile (left) defines the exact match length at each position along the query with respect to the reference. Using a “shredded” sampled document array, matches are mapped back to reference regions to identify a cluster of matches. Here, a read maps to Ref 5, which is the predicted reference (in red).

**Figure 2:**
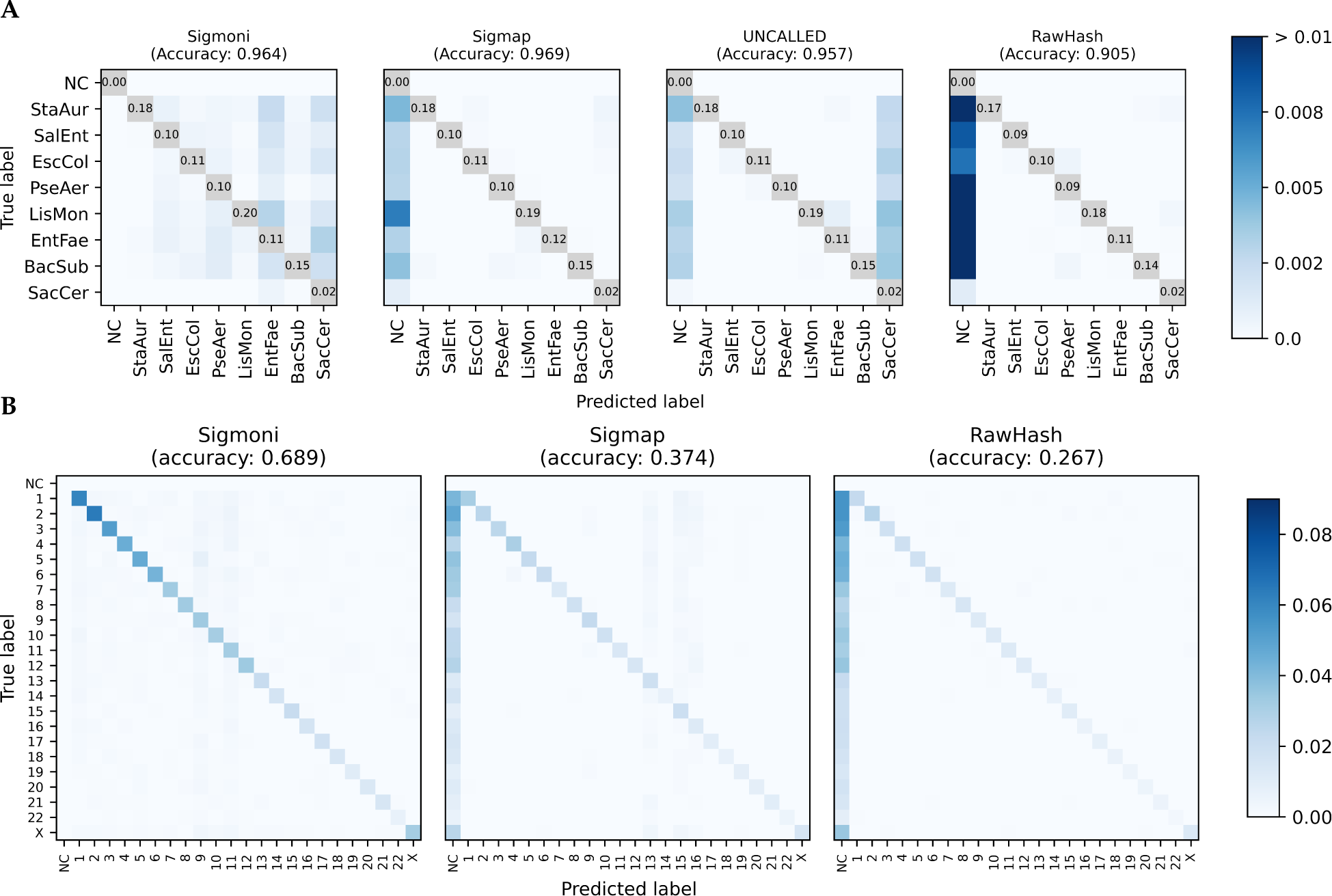
**(A)** Comparison of of mock community multi-class classification confusion matrices for each signal-based method. The diagonal (TP) is omitted, with proportion of reads provided instead to high-light off-diagonal (misclassified) reads. **(B)** Confusion matrix of human chromosome-level classification of NA12878 reads against CHM13. As the donor individual is female, ChrY was omitted from the reference. NC = not classified.

#### Human chromosome

As a more challenging scenario, we also tested the methods’ ability to distinguish reads from different human chromosomes, as might be needed in an experiment targeting particular chromosomes [23]. We queried reads from NA12878 against the CHM13 reference genome, omitting the Y chromosome since the donor individual is female. We classified reads by chromosome, including autosomes and the X chromosome. Similar to the Zymo reads, true chromosomal-origin labels were computed by querying basecalled reads against the NA12878 reference genome (to avoid reference bias) using Minimap2. For this experiment, RawHash was run in its -x faster mode, as is suggested for references longer than 3 Gbp. UNCALLED was not able to build an index over the human chromosomes.

Sigmoni was able to classify significantly more reads than Sigmap and RawHash; Sigmap did not classify ∼62% and RawHash did not classify ∼74% of the NA12878-origin reads (Figure 2**B**). Sigmoni also out-performed both other tools in accuracy: 0.689 for Sigmoni versus 0.374 for Sigmap and 0.267 for RawHash.

### 2.4 Adaptive sampling

Nanopore sequencers from Oxford Nanopore can selectively eject DNA molecules from pores, a facility called “adaptive sampling.” For greatest effect, the ejection decision should come within a few seconds of the molecule entering the pore. To compare these methods based on their utility for adaptive sampling, we ran each on increasing-length prefixes of the signal, simulating a scenario where each tool was run on a sequence of signal “chunks” in real-time. A typical chunk consists of about 4,000 picoamp measurements, or about ∼420bp after basecalling. Classifying reads efficiently and accurately on fewer chunks yields greater time/cost savings during sequencing.

We performed the same binary classification tasks as described in Results 2.2, identifying yeast reads from the bacterial components of the Zymo mock community, as well as human reads versus the entire Zymo mock community. We compared tools using the F1 metric without any length-weighting, since the signal chunks were already the same length for each read. Lastly, we evaluated classification efficiency for each tool for both tasks as well as the classification rate. Timing results were performed using a single thread of execution for each tool, on a 3 GHz Intel Xeon Gold Cascade Lake 6248R CPU with 1.5TB DDR4 2933MHz memory.

In the mock community experiment, Sigmap had the highest F1, particularly over the first few chunks (Figure 3A). RawHash had similar but slightly lower F1. UNCALLED had high F1 on a single chunk, but fell behind both Sigmap and RawHash for larger numbers of chunks. While Sigmoni had clearly lower F1 for small numbers of chunks, it gradually improved its F1 to greater than UNCALLED’s after 4 chunks, and comparably to Sigmap’s and RawHash’s after 10 chunks.

**Figure 3:**
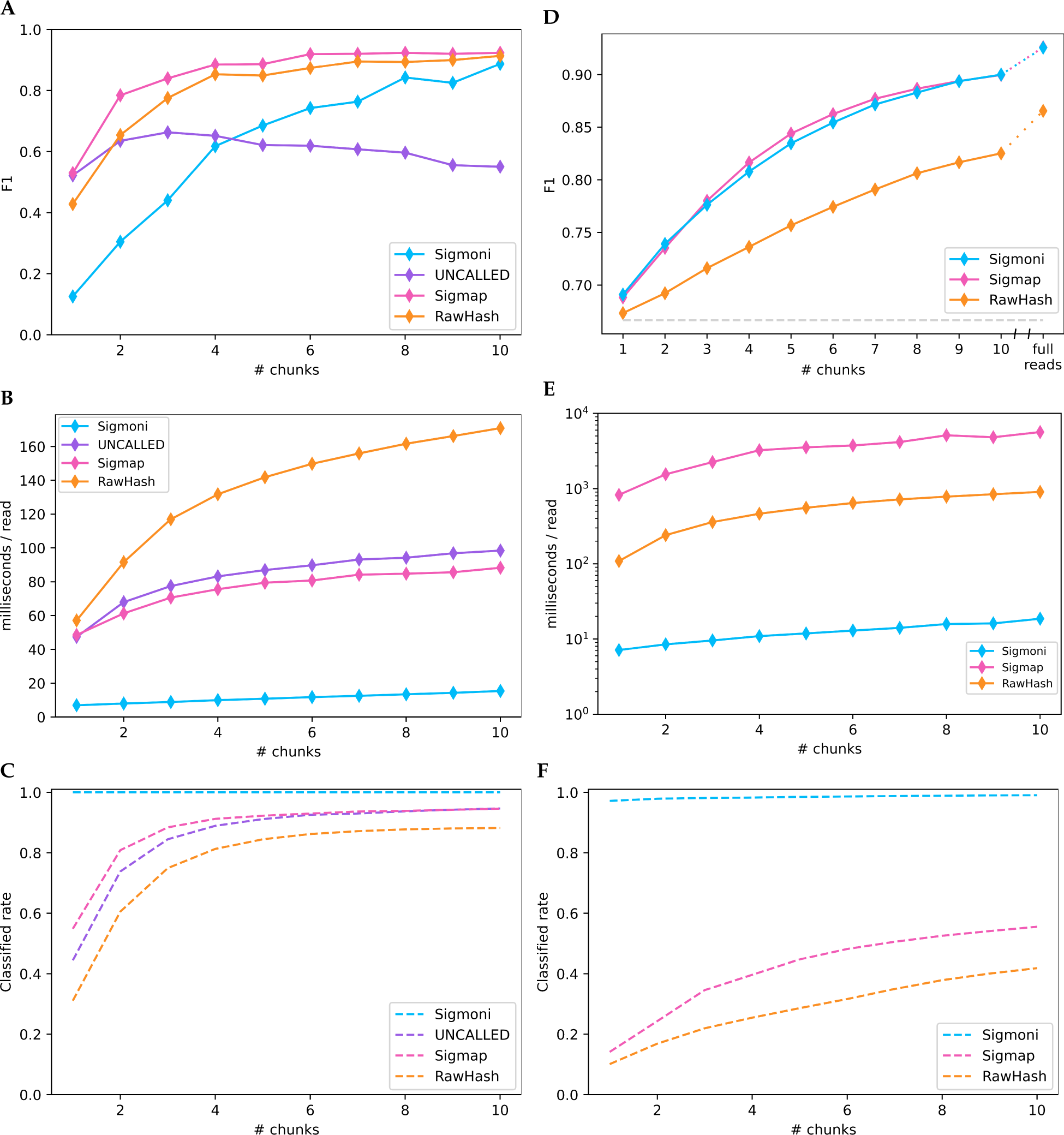
**(A-C)** Binary classification of yeast-origin reads from a mock community on “chunks” of signal. Each chunk represents 1 second of sequencing, ∼420bp; **(A)** F1 score (unclassified reads are considered bacterial in origin), **(B)** classification speed for increasing length signal chunks, **(C)** proportion of reads classified by each method. **(D-F)** Binary classification on a hybrid dataset of human-origin (NA12878) reads and Zymo mock community reads; **(D)** F1 score (unclassified reads are considered human-origin, as in the case of a “host depletion” experiment), **(E)** classification speed, **(F)** read classification rate.

Sigmoni was much faster than all competing methods (Figure 3B), 7× faster than Sigmap and UN-CALLED and 8× faster than RawHash on a single chunk. For Sigmoni, classification time grew linearly with respect to query length. The time taken by the competing tools grew sublinearly, possibly due to their use of a more complex seed-chain-extend mapping strategy.

For the host depletion task, Sigmoni achieved similar F1 as Sigmap (Figure 3D), while both tools had much higher F1 than RawHash. Sigmoni was significantly faster than both tools (Figure 3E), between 100– 300× faster than Sigmap over the first ten sequencing chunks and between 15–50× faster than RawHash (again run using the recommended -x faster setting). On certain read lengths, Sigmap took longer than a second to classify a read on average, exceeding the time needed to sequence the chunk itself.

On a single chunk, both RawHash and Sigmap are unable to classify more than 80% of reads, while Sigmoni can classify over 97% of reads from the first chunk with a similar accuracy (Figure 3F). In a host depletion scenario, this may be acceptable since unclassified reads can be assumed to be human-origin. But Sigmoni’s ability to make a decision for each read makes it particularly applicable in applications where high sensitivity is needed and the reference is large.

### 2.5 Pangenome scaling

An advantage of the compressed *r*-index is how it scales to large, repetitive pangenome references. Unlike the FM-index, which scales linearly with reference length, the *r*-index scales proportionally to *r*, the number of runs in the Burrows-Wheeler Transform (BWT). The BWT rearranges the letters of the input according to the alphabetical order of their right contexts, so more repeitive inputs yield fewer and longer runs. As such, adding highly similar genomes to the reference will increase index size sublinearly. SPUMONI 2 [16] highlights this sublinear growth for growing collections of *E. coli* genomes.

We found that *r*-index’s scaling is similarly sublinear with respect to quantized nanopore signal (which we call “bin sequences”). That is, when computed over the binned pangenome, *r* still grows quite sublinearly, leading to small indexes. Table 2 compares the size of the signal-space *r*-index when used to index various numbers of human genome assemblies, including from the Human Pangenome Reference Consortium (HPRC) [24]. Notably, an *r*-index over the set of binned HPRC genome assemblies fit in ∼13 GB. As a comparison, we built a RawHash index (the only other signal-based tool that could index the HPRC assemblies), which used 1,082 GB.

**Table 2:**
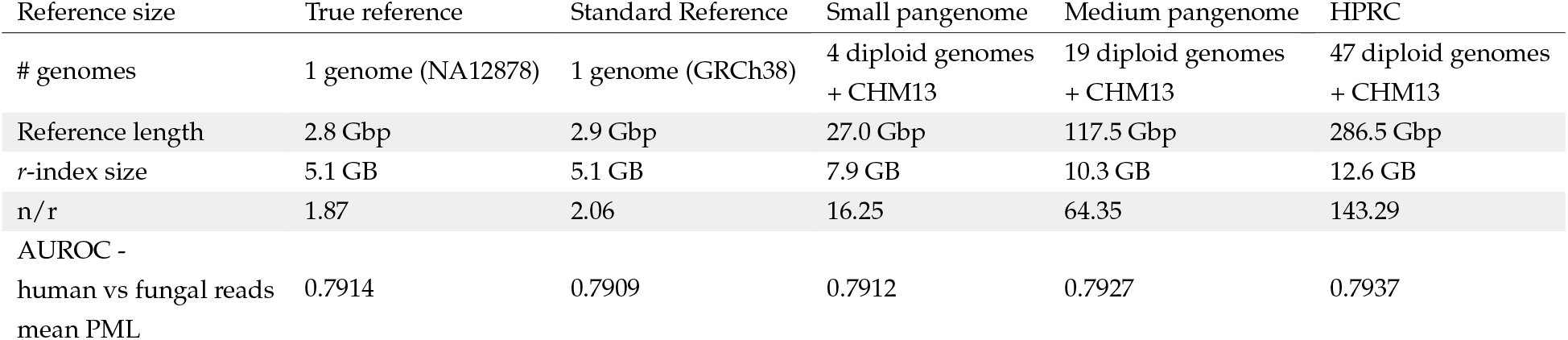
Growth of *r*-index with increasing sets of human reference assemblies. Reads from NA12878 were queried against larger references and the exact matching lengths (PMLs) were compared to a null distribution, yeast-origin reads queried to human references. The area under the receiver operator characteristic curve (AUROC) metric measures the discernibility of PML distributions between the two read query sets.

The *n/r* ratio relates *n*, the number of picoamp alphabet characters in the reference text *T*, to *r*, the number of runs in BWT(*T*). As assemblies are added to the pangenome, *n* increases by roughly 3 Gbp for each assembly whereas *r* increases marginally since most of the newly added sequence is redundant.

To evaluate how including more reference assemblies affects classification, we queried two sets of reads simulated from the NA12878 reference and yeast (*S. cerevisiae*) genome, against 5 references: a personalized diploid reference for NA12878, the standard human genome reference GRCh38, 4 diploid assemblies from HPRC + CHM13, 19 diploid assemblies from HPRC + CHM13, and the entire HPRC set consisting of 47 diploid assemblies + CHM13. Reads were simulated using Squigulator [25], with mean read length of 10,000bp, amplitude domain noise factor of 2.0, and dwell time standard deviation of 8.0. We compared the average pseudomatching lengths (PMLs) from NA12878 reads vs. yeast-origin reads, and computed the area under the receiver operator characteristic curve (AUROC) to measure how well the PMLs distinguished the classes. We observed that adding more assemblies increased the AUROC, highlighting the utility of pangenomes in overcoming reference bias and improving classification.

## 3 Discussion

Sigmoni adapts the *r*-index classification framework to analysis of nanopore signal data using a combination of picoamp binning, a sampled document array structure for computing co-linearity statistics, and a novel classification method that accurately classifies reads using easy-to-compute pseudo-matching lengths. By avoiding the complexities of the seed-chain-extend paradigm, Sigmoni’s core algorithm consists only of a simple linear-time loop. Besides its efficiency, the algorithm is also easy to pause and resume as more data becomes available, since very little “state” is maintained across loop iterations.

Sigmoni also highlights the applicability of the *r*-index to arbitrary alphabets. By binning picoamp space into a small discrete alphabet, Sigmoni can also adapt to newer Nanopore chemistries (like R10), where larger *k*-mers affect the signal output. This is in contrast to *k*-mer based methods, where the *k* parameter significantly affects runtime and index size. By avoiding the parameters inherent to other methods — e.g. *k*-mer length, seed-chain-extend parameters — Sigmoni can avoid over-fitting to a specific classification task.

Previous studies have used lightweight basecallers for faster adaptive sampling [13]. Our picoamp binning procedure could be considered a very simple “basecalling” step. Rather than simply binning, we could have implemented more sophisticated methods used in basecallers, such as Hidden Markov Models (HMMs). Indeed, any method that preprocesses the picoamp signal into a sequence of discrete alphabet characters — whether they represent picoamp ranges, HMM states, or nucleotides — could fit in the Sigmoni and *r*-index frameworks, which can handle arbitrary-size discrete alphabets.

Other methods introduced here are also applicable to read classification in the context of basecalled reads. For instance, document shredding (4.4) could be used to increase location resolution for nucleotide-space exact matches. These methods can also aid in detecting large-scale structural genomic variation, which can be more difficult to represent using graph-based approaches.

Currently Sigmoni requires both a positive and null reference example to map reads against. However, in certain scenarios, the “null” reference might be unknown or simply any sequence not contained within the positive reference. SPUMONI achieves this using null statistics, however this null-based comparison is difficult with noisy signal data. As future work, we are exploring generative HMMs which can create realistic “null” reference bin sequences that maintain a similar sequence composition, while serving as a point of comparison for matching lengths to determine whether a read belongs to the positive reference database. This idea could also benefit basecalled classification methods like SPUMONI.

## 4 Methods

### 4.1 Event Detection and Normalization

The input to Sigmoni is the sequence of regular picoamp measurements from the pore. Sigmoni uses the Uncalled4 software (https://github.com/skovaka/UNCALLED/tree/uncalled4) to perform event detection, grouping stretches of picoamp (*pA*) measurements into “events” that correspond roughly to the traversal of a single nucleotide through the pore. Uncalled4’s event detection algorithm is adapted from Scrappie (https://github.com/nanoporetech/scrappie), performing a rolling *t*-test to find significant changes in picoamp level. Sigmoni defaults to the event detection parameters proposed in the Sigmap study [7], which are designed to balance stay and skip errors. Sigmoni normalizes the picoamp values observed to match those encoded in the pore model, which defines the expected current in *pA* for each *k*-mer. Here, Sigmoni uses the same normalization algorithm as UNCALLED [5], based on the Welford algorithm.

### 4.2 Binning

#### Pore model

Sigmoni uses a pore model for normalization and signal binning. A pore model maps each *k*-mer to the expected picoamp value produced by passing the *k*-mer sequence through a nanopore. The value of *k* is determined by the number of nucleotides within the nanopore which affect the current output, which depends on pore chemistry. Sigmoni uses this expected *pA* model to convert the reference sequence into expected current signal for binning.

#### Defining bins

Given the normalized event sequence, Sigmoni next discretizes the continuous current values into an alphabet of picoamp ranges. Using wider (and fewer) picoamp ranges allows for greater robustness to noise in the signal. Let the picoamp (*pA*) width of each bin be *s*_*p*_, which is determined by dividing the range of the expected *pA* values from the pore model, [min_*p*_, max_*p*_], by the chosen number of bins *n*. For an event *e*, defined by its mean current and dwell time (*e*_*c*_, *e*_*t*_), the corresponding bin *b* is computed using equation 1.

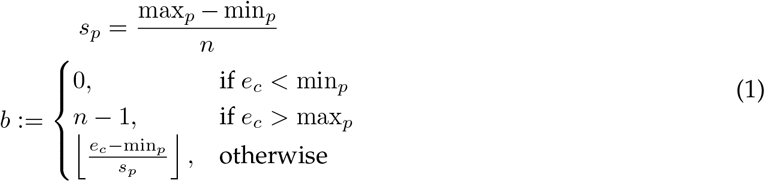

#### Reference Binning

Sigmoni first iterates over the *k*-mers of the reference sequences in order. Using a pore model, it converts *k*-mers to the expected current value in *pA*. The sequence of *pA*s is binned using the same procedure as above. As described below, both the read and reference are also homopolymer compressed. This compressed sequence of *pA* alphabet characters serves as the new reference, which is written to a FASTA file using printable ASCII characters to represent the alphabet, and is indexed by *r*-index.

#### Homopolymer compression

Ideally, the event detection algorithm creates one event per nucleotide. In practice, errors may occur: (1) *stays* result when two or more consecutive events correspond to a single *k*-mer (over-segmented), and (2) *skips* occur when a single event spans multiple *k*-mers. Skips are difficult to recover from after-the-fact since the resulting signal convolves multiple *k*-mers. Stays have a recognizable signature, appearing as consecutive events with similar *pA* values, which may in turn appear as runs of identical picoamp-alphabet characters. Sigmoni handles stay errors by compressing same-character runs as a single character, similar to homopolymer compression (HPC) of nucleotide sequences. The reference sequence is compressed in the same way, ensuring that true repeats in the reference are still queryable.

### 4.3 Compressed index and matching statistics

The *r*-index, proposed by Gagie et al. [26], [27], consists of a run-length encoded Burrows-Wheeler Transform (RLEBWT), as well as auxiliary data structures that allow for querying of match locations in *O*(*r*) space. MONI [20] introduced an additional auxiliary “thresholds” structure together with a two-pass algorithm for computing matching statistics (MS) with respect to the enhanced index. The MS vector, which is the same length as the query string, defines the length of the longest exact match at each position of the query to the reference that are non-extendable to the right (i.e. half maximal exact matches, half-MEMs). SPUMONI [4] modified this method to instead compute pseudo-matching lengths (PMLs), a kind of truncated matching statistic that can be computed in a single-pass. The simple loop for computing PMLs makes this algorithm particuarly well suited to applications requiring real-time decisions like adaptive sampling. Also, computing PMLs requires fewer of the auxiliary data structures needed by MONI (e.g. the sampled suffix array), yielding a smaller index. Sigmoni uses the SPUMONI software to build the *r*-index over the binned and homopolymer compressed reference sequence and computes PMLs along the binned query signal. Large PMLs represent stretches of query signal that match the expected current of a corresponding reference region.

### 4.4 Document array

A drawback to *r*-index PML computation is the inability to compute match location information efficiently, since neither the suffix array nor any sampled version of it is stored in the index. To address this, SPUMONI 2 [16] introduces a “sampled document array” structure. Suppose the reference consists of a collection of sequences (“documents”); the sampled document array stores a document label at the start and end of each run in the RLEBWT according to the document of origin for the suffix at that position. During PML computation, matches can be labeled with document labels encountered while traversing the BWT. This data structure increases the overall index size (though it is still O(*r*) altogether), while enabling multiclass classification of reads based on the document labels found for long matches.

This enables only document-level locality information (such as genomes in a pangenome). Sigmoni extends the idea to allow for higher-resolution locality information by “shredding” the documents. Each reference sequence is partitioned into equal-length, non-overlapping substrings called “shreds.” Each shred is assigned a distinct document label. The document array is built using SPUMONI and reads are queried in the same way, however the document labels for PMLs now correspond to shreds within the larger pangenome, indicating a specific local region of the reference where the match occurs. By default, Sigmoni uses a shred size of 100 kbp.

The original document array takes *O*(*r* log *c*) space, where *c* is the number of documents. With equal-sized shreds, the number of documents increases linearly with reference length, i.e. *c ∈ O*(*n*), and the full shredded document array takes *O*(*r* log *n*) space. Though this structure is no longer strictly *O*(*r*), setting the length of shreds to be large enough (*>* 100 kbp) enables feasible data structure size in practice.

### 4.5 Classification

#### Weighted Document Vote

To classify a read, Sigmoni uses a plurality vote of document labels, weighted by the corresponding PML lengths, at “peak” PMLs. Specifically, for a read of length *n* with PML vector *P* and document vector *D*, Sigmoni uses the rule defined in equation 2,

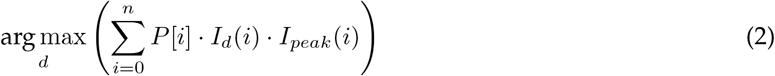

where *I*_*d*_(*i*) is the indicator function for when *D*[*i*] = *d*, and *I*_*peak*_(*i*) is the indicator function for “peak” PML positions, defined as any position *i* such that *P* [*i*] ≥ *P*[*i* − 1]. Since PMLs decrement along the query within a single distinct match, any jump in PML represents a new match. The peak PML of a match is used to eliminate over-representation of long matches.

Though each PML is labelled with a single document ID, as pointed out in SPUMONI 2 [16], the match may exist in multiple documents. However, longer matches tend to be more unique (i.e. the exact match appears in a single document), therefore by taking a weighted vote, we can infer the true document of origin for the full read. Lastly, using shredded documents (Methods 4.4) avoids the issue of over-representation of matches to longer documents (as each shred is equal-sized), preventing classification bias in both multiclass and binary tasks.

#### Binary classification

Sigmoni can perform binary classification in cases where sequences representing the two classes are both present in the reference index. Here we assume that the two classes represent a “positive” class containing on-target sequences of interest and a “null” class containing off-target or random sequences. In this scenario, Sigmoni will classify reads in a similar manner to multiclass classification, but rather than choose the document with the highest weighted sum of PMLs (largest “spike”), the top documents from both positive and null references are compared. The read is classified as originating from the positive class if the weighted sum for the top positive-class document is sufficiently larger than that of the null-class document. The “sufficiently larger” criterion is judged by testing if the ratio is greater than some threshold (“spike ratio threshold”). The threshold can be computed empirically during a ”burn-in” period at the start of the sequencing process, and should correlate with the expected ratio of positive to null class reads. That is, the task of identifying reads from a relatively low-abundance positive class should use a higher threshold. Removing the threshold (and choosing the top document with equation 2) is suitable for scenarios where classes are present in roughly equal proportion.

#### Sequence complexity correction

Large PMLs can result from matches to low-complexity regions in the references, such as tandem repeats. Long matches of this kind are more likely to happen by coincidence and are less likely to represent the true origin of the read. To account for this, we scale terms in equation 2 by a factor *C*_*d*_, which should equal the sequence complexity for document *d*. This factor can be estimated by computing a measure of sequence complexity like empirical entropy or substring complexity *δ* [28] over the concatenation of exact matches, or alternatively during index building for each reference shred.

## 5 Acknowledgments

This work was carried out at the Advanced Research Computing at Hopkins (ARCH) core facility, supported by the National Science Foundation (NSF) grant number OAC 1920103. We thank Michael Schatz for his advice and comments on the manuscript.

## 7 Availability of data and materials

Sigmoni is an open-source software available at https://github.com/vshiv18/sigmoni. For the Zymo mock community dataset, we used reads available under SRA accession SRX7711546 from Kovaka et al. [5]. For NA12878 reads, we used reads from the NA12878 WGS consortium [22], specifically from flowcell FAB42260 under the rel6 data. Genome assemblies from HPRC [24] were obtained from https://github.com/human-pangenomics/HPP_Year1_Assemblies/blob/main/assembly_index/Year1_assemblies_v2_genbank.index. Bacterial reference genomes for the mock community were obtained from ZymoBIOMICS (https://s3.amazonaws.com/zymo-files/BioPool/D6322.refseq.zip), and the RefSeq S288C assembly was used for *S. cerevisiae* (accession GCF 000146045.2). The NA12878 reference genome was also used (accession GCA 002077035.3) for pangenome analysis and read simulation.

## 8 Authors’ contributions

VSS and BL designed the method, with help from OYA, SK, and MZ. VSS wrote the software and performed the experiments. All authors contributed to the manuscript.

